# Autism associated mutation in *Cacna1d* causes perseverative behavioral phenotype in mice

**DOI:** 10.64898/2026.01.13.699343

**Authors:** Shintaro Otsuka, Jian Xu, Anis Contractor

**Affiliations:** Department of Neuroscience, Northwestern University Feinberg School of Medicine, Chicago, IL 60613

## Abstract

Behavioral inflexibility and perseveration are core features of autism spectrum disorder and are frequently modeled in mice using reversal learning and repetitive behavior assays. Mutations in *CACNA1D*, which encodes the L-type calcium channel Cav1.3, have been linked to autism, yet their behavioral consequences remain incompletely characterized. We examined mice carrying the autism-associated *Cacna1d*^G407R^ gain-of-function mutation across a battery of assays assessing learning, flexibility, and repetitive behavior. Mutant mice learned spatial discriminations and instrumental contingencies at rates comparable to wild-type controls but exhibited deficits during reversal learning following intermediate and extended overtraining, as well as under probabilistic reinforcement. Across ethological assays, mutant mice showed increased grooming, marble burying, and nestlet shredding, consistent with enhanced perseverative behavior. Anxiety-related measures and general locomotion were largely unaffected.

These results identify Cav1.3 gain-of-function as a selective regulator of behavioral flexibility and support a role for calcium-dependent corticostriatal plasticity in autism-associated perseveration.

## Introduction

Autism spectrum disorders (ASDs) are a group of highly prevalent neurodevelopmental conditions characterized by deficits in social communication, restricted interests, and repetitive or perseverative behaviors (Lei et al., 2022). ASDs have a strong genetic component and large-scale sequencing studies have revealed numerous rare pathogenic *de novo* variants in genes encoding synaptic and ion channel proteins (De Rubeis et al., 2014; Iossifov et al., 2012), implicating alterations in neuronal excitability and plasticity as key mechanisms underlying autistic behaviors. Despite the many advances in genetic studies of autism (Iossifov et al., 2014; Satterstrom et al., 2020), the functional impact of many autism-associated mutations on neural circuits and behavior remains poorly understood.

Mutations in genes encoding L-type voltage-gated Ca²⁺ channels have emerged as recurrent causes of neurodevelopmental disorders (Gargus, 2009; Pinggera et al., 2015). The Cav1.2 and Cav1.3 channels, encoded by CACNA1C and CACNA1D, respectively, regulate dendritic Ca²⁺ entry that couples neuronal activity to transcriptional and synaptic plasticity processes (Deisseroth et al., 1998; Mermelstein et al., 2000; Zhai et al., 2024). Multiple *de novo* CACNA1D missense variants have been identified in individuals with autism (Iossifov et al., 2012; O’Roak et al., 2012; Pinggera et al., 2015; Pinggera et al., 2017; Hofer et al., 2020). The *de novo* CACNA1D p.G407R variant was identified in a male individual (first symptomatic at ∼15 yrs) with an autism spectrum disorder diagnosis, and at the time of reporting had no documented intellectual disability, no seizures, and no overt endocrine or neurological comorbidities (Pinggera et al., 2015; Ortner et al., 2020). Functionally the variant produces a strong effect on gating of the Cav1.3 channel, shifting channel activation to more hyperpolarized potentials and impairing inactivation, producing a gain-of-function phenotype with enhanced Ca²⁺ influx (Pinggera et al., 2015). Such changes are predicted to alter neuronal excitability and synaptic development in brain regions where Cav1.3 is strongly expressed. One such region is the striatum which is a major site of Cav1.3 expression and is critically involved in action selection, habit formation, and behavioral flexibility (Graybiel, 2008). Aberrant corticostriatal signaling has been implicated in several core symptoms of autism, including repetitive and perseverative behaviors (Burguiere et al., 2015). Striatal spiny projection neurons (SPNs) depend on Cav1.3 channels for dendritic Ca²⁺ signaling, synaptic plasticity, and spine maintenance (Day et al., 2006). Moreover, Cav1.3 channels interact with the synaptic scaffold SHANK3 (Zhang et al., 2005), a synaptic protein with established association to ASD (Soorya et al., 2013; Satterstrom et al., 2020), suggesting that perturbation of Cav1.3 function could directly impact corticostriatal synaptic function. Despite these observations, the circuit and behavioral consequences of autism-associated gain-of-function mutations in Cav1.3 remain largely unexplored.

Perseverative and repetitive behaviors are among the most robust and defining behavioral features of ASD. These behaviors range from simple motor stereotypies to complex cognitive inflexibility, such as difficulty shifting strategies or adapting to changing contingencies (Ragozzino, 2007; Miller et al., 2015). In human neuroimaging studies abnormal striatal activation and connectivity are observed in individuals with autism during tasks requiring reversal learning or flexible decision-making (D’Cruz et al., 2016). Similarly, rodent models of autism show impairments in operant reversal learning, increased habit formation, and reduced behavioral flexibility (Amodeo et al., 2012; Copping et al., 2017), indicating that disrupted striatal function may underlie perseverative responding across species. Thus, assessing reversal learning and behavioral flexibility in genetic models of autism provides a powerful translational approach to identify circuit-level mechanisms contributing to core ASD-related behaviors.

We recently generated a knock-in mouse model carrying the *Cacna1d*^G407R^ mutation identified in individuals with autism (Zhai et al., 2024). This mutation enhances the activity of Cav1.3 L-type calcium channels, leading to a gain-of-function increase in calcium influx which we measured in striatal projection neurons in these mice.

Moreover, we found that synaptic plasticity of corticostriatal synapses was altered in the mice through a postsynaptic mechanism that is dependent upon L-type Ca2+ channels (Zhai et al., 2024). Building on this foundation, in the present study we examined behavioral phenotypes in the *Cacna1d*^G407R^ mice examining measures of behavioral inflexibility and perseverative responding as well as social behaviors

## Results

### Spontaneous and induced behaviors

*Cacna1d*^G407R^ heterozygous knock-in mice were examined in several common behavioral paradigms alongside wildtype littermates. We first tracked mice in an open field test (OFT) to assess spontaneous locomotor activity by measuring total distance traveled, providing a baseline measure of general activity in both male and female mice (Xu et al., 2017). Additionally, analyzing distance traveled separately for males and females allowed us to account for potential sex-dependent differences in baseline activity levels. We found that for female mice, distance travelled declined across time bins (main effect of time; LR = 111.4, df = 18, p < 1.0 × 10⁻¹ ), indicating habituation, but there was no overall genotype effect (LR = 9.72, df = 10, p = 0.47) and no genotype × time interaction (LR = 8.47, df = 9, p = 0.49), indicating that locomotor activity did not differ between female WT and *Cacna1d*^G407R^ mice, as assessed with a linear mixed-effects model (random intercept for subject) (**Figure 1A**). Similarly for male mice analysis of locomotor activity in the open field demonstrated that distance decreased significantly over the 30-minute session (main effect of time: p < 0.001), reflecting normal habituation. There was no significant effect of genotype (p = 0.536) and no significant genotype × time interaction (p = 0.134), indicating that WT and mutant males exhibited comparable locomotor activity and habituation patterns (linear mixed-effects model with distance traveled as the dependent variable, genotype as a fixed factor, time bins as repeated measures, and mouse ID as a random effect) (**Figure 1B**).

**Fig. 1.**
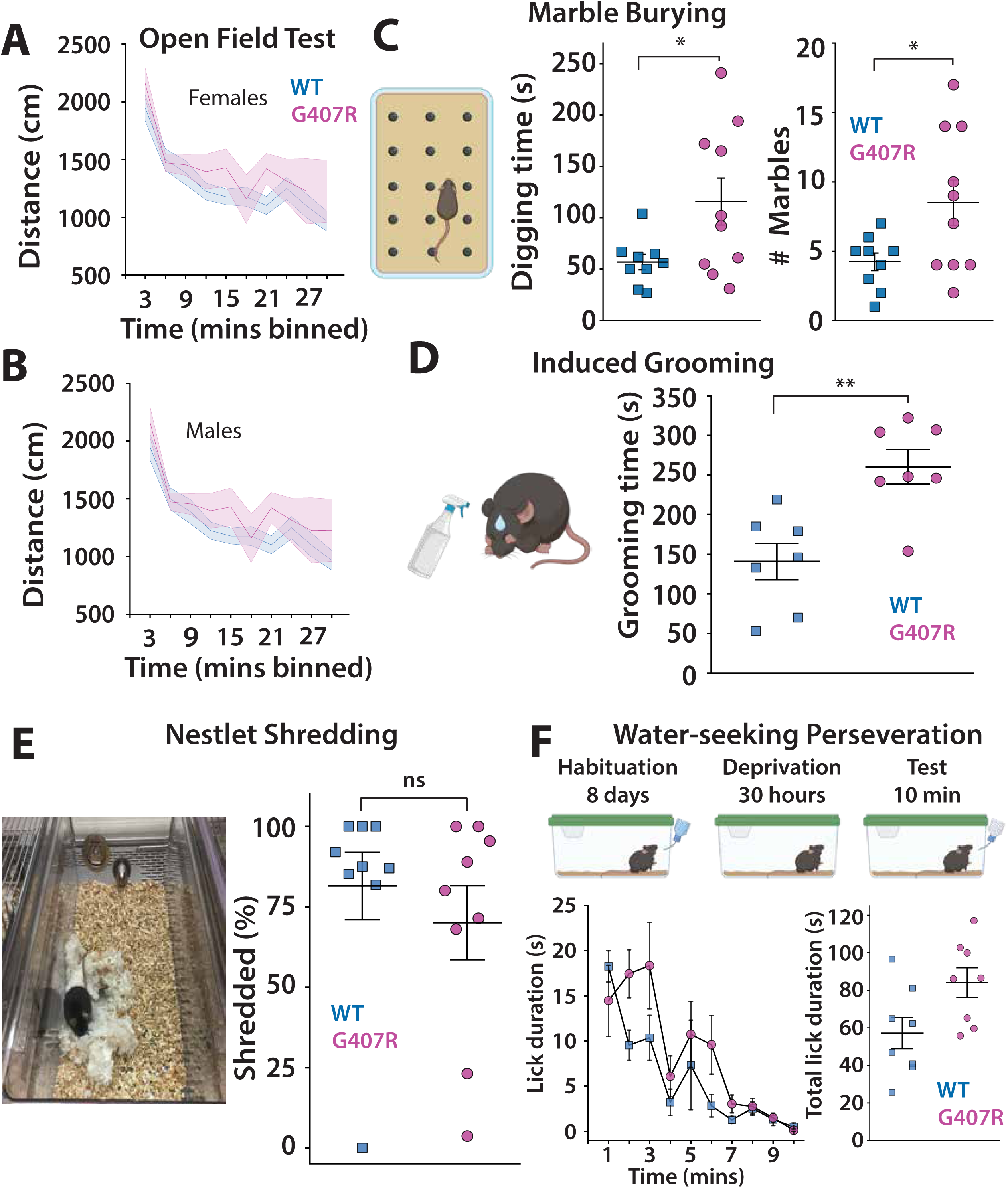
*Cacna1d*^G407R^ mutant mice demonstrate enhanced repetitive behavior. (**A**, **B**) Locomotor activity of female (**A**) and male (**B**) *Cacna1d*^G407R^ heterozygous mice and wild-type controls in the open field test. Females: *n* = 7 (wild-type) and *n* = 5 (heterozygous); males: *n* = 10 (wild-type) and *n* = 5 (heterozygous). (**C**) Schematic of the marble-burying task (left). Results of the marble burying test (Right). *Cacna1d*^G407R^ heterozygous mice exhibited prolonged digging time (left panel) and increased marble burying behavior (right panel). *n* = 9 (wild-type) and *n* = 10 (heterozygous). (**D**) Schematic of the induced grooming task (left). *Cacna1d*^G407R^ heterozygous mice displayed increased grooming behavior. *n* = 7 (right panel). (**E**) Representative example of nestlet shredding (left panel). The ratio of nestlet shredding did not differ between genotypes. *n* = 9 (right panel). (**F**) Top panel show schematic of Water-seeking perseveration task. Timecourse of licking during the test (1min bins) (Bottom left panel) Total lick duration during during 10 min test (bottom right panel) *n* = 8 (wild-type) and *n* = 8 (heterozygous). Schematics were generated in bioRender.

Next we assessed repetitive, perseverative digging using the marble-burying assay, a widely used paradigm that quantifies species-typical digging/burying behaviors and is commonly interpreted as a measure of compulsive and repetitive-like behavior rather than simple novelty-induced anxiety (Thomas et al., 2009; Angoa-Perez et al., 2013). Mice were introduced into a new cage containing 24 marbles placed equidistance apart on top of 3-5cm of bedding (Xu et al., 2017). Digging time and number of marbles buried were measured during the 15-minute test in both male and female mice as we had done previously (Xu et al., 2017). Digging time was significantly different between WT and *Cacna1d*^G407R^ heterozygous mice showing elevated digging (Welch’s t-test: t = –2.45, p = 0.032). Marble burying behavior was also assessed by counting the number of marbles fully or partially covered by bedding. Because the two groups differed in variance, we again used Welch’s t-test to compare genotypes. This analysis revealed a significant difference between WT and heterozygous mice (t = –2.44, p = 0.032), with *Cacna1d*^G407R^ heterozygous mice burying a greater number of marbles (**Figure 1D**).

In order to assess self-oriented perseverative behaviors beyond digging and marble-burying, we measured grooming behavior following a light water mist applied to the forehead of each mouse. Immediately after the spray, each animal was placed in a clean testing chamber, and the duration of self-grooming was measured over a ten-minute period. This spray-evoked grooming paradigm has been previously used as an index of maintenance or stereotyped behavioral responses to assess repetitive or perseverative behaviors in models of neurodevelopmental disorders (Moretti et al., 2005; Braz et al., 2015). In our cohort, water spray evoked grooming duration differed significantly between genotypes, with *Cacna1d*^G407R^ heterozygous mice grooming longer than WT mice (p = 0.0027, Welch’s t-test) suggesting an enhanced maintenance or perseverative response in the heterozygous genotype. This result is consistent with the notion that elevated grooming after a mild perturbation (such as water mist) may reflect an increased propensity for repetitive self-maintenance or perseverative behaviors, rather than merely a transient stress reaction.

We also assessed nestlet-shredding behavior as an additional measure of repetitive or self-directed activity. In this assay, each mouse was provided a single pre-weighed nestlet of cotton fiber in a standard cage for 12 hours during the dark cycle, after which the remaining intact material was weighed and the amount shredded calculated. The nestlet-shredding test has been used in rodent models of compulsive and perseverative behavior but also interpreted as an index for engagement with novel objects (Dorninger et al., 2020). Originally developed in the context of obsessive-compulsive disorder and autism spectrum models to complement the marble-burying test, studies interpret increased shredding as reflecting excessive repetitive or maintenance-oriented behavior (Angoa-Perez et al., 2013). We quantified the percentage of a standard cotton nestlet shredded by each mouse over the testing period. WT mice shredded 81.50 ± 10.45%, n = 9 of the nestlet, whereas G407R mutant mice shredded 70.06 ± 11.49%, n = 9. Comparison using a non-parametric Mann–Whitney U (both groups violated normality assumptions; Shapiro–Wilk p < 0.05), analysis revealed no significant difference between genotypes (U = 50, p = 0.42). Thus, unlike other repetitive behaviors such as digging, grooming, and marble burying, nestlet shredding was not different between WT and mutant mice. Considering the converging profile of increased digging time and grooming duration these data support an interpretation of the mutant mice as exhibiting an elevated repetitive-behavior phenotype rather than an isolated alteration in exploration or anxiety.

To further examine how the *Cacna1d*^G407R^ mutation affects acquired behaviors, we developed a novel assay termed the *Water-Seeking Perseveration Task* (**Figure 1** **F**). This task assesses behavioral flexibility by measuring the persistence of licking behavior after reinforcement is withdrawn. Lick duration was quantified in 1-min bins over a 10-min probe period after an empty water bottle was returned to the home cage. A two-way repeated-measures ANOVA revealed a significant main effect of time, indicating progressive extinction of licking behavior across the session (RM-ANOVA, main effect of time, p < 0.001), as well as a significant main effect of genotype (RM-ANOVA, main effect of genotype, p < 0.05). Importantly, there was a significant genotype × time interaction (RM-ANOVA, interaction, p < 0.05), reflecting a failure of *Cacna1d*^G407R^ mice to appropriately suppress licking over time (**Figure 1F**). Post hoc comparisons confirmed that mutant mice exhibited significantly elevated licking during later minutes of the session relative to wild-type controls (Sidak-corrected multiple comparisons, p < 0.05), consistent with enhanced perseverative responding.

Furthermore, total lick duration across the 10-min session was calculated for each animal. *Cacna1d*^G407R^ mice exhibited significantly greater total licking compared with wild-type littermates (Mann–Whitney U test, p < 0.05), indicating a sustained inability to disengage from a previously reinforced behavior (**Figure 1F**). Together, these findings demonstrate that the *Cacna1d*^G407R^ mutation impairs behavioral flexibility across ethologically relevant contexts, reflected by persistent engagement in a previously reinforced action despite a change in environmental contingencies.

### Tests of anxiety

To directly address whether mutant mice demonstrated changes in anxiety we adopted several standard paradigms to compare the behaviors of genotypes. In our analysis of the open field test (OFT) we had measured time mice spent in the center area of the arena to assess whether genotype influenced anxiety-related exploratory behavior. We quantified the total distance each animal traveled within the center region of the open-field arena. WT mice traveled 624.10 ± 22.52 cm, n = 17 mice whereas G407R mutant mice traveled 575.21 ± 41.39 cm, n = 9. Both groups were normally distributed and exhibited comparable variance, supporting the use of a parametric test. A Welch’s t-test revealed no significant difference in center distance between genotypes (p = 0.32).

Thus, despite the pronounced differences observed in repetitive behaviors such as digging and grooming, center exploration, an index of anxiety-like behavior, did not differ between WT and mutant mice, suggesting that the enhanced repetitive behaviors observed in the mutant group are unlikely to be secondary to altered anxiety levels.

As further measures of anxiety, we tested mice in the elevated zero maze, a circular variant of the elevated plus maze that provides a continuous path and eliminates the center decision zone (**Figure 2A**). This assay is widely used to quantify anxiety-like avoidance behavior in rodents, with increased exploration of the open, unprotected quadrants interpreted as reduced anxiety and decreased open-zone exploration interpreted as heightened anxiety (Shepherd et al., 1994). Mice freely explored the maze for 10 minutes and were tracked by overhead video. Across all measures of zero-maze behavior that we made including time spent in each quadrant (**Figure 2B**), distance traveled in each quadrant (**Figure 2C**), and quadrant entries (**Figure 2D**), WT and G407R mutant mice showed comparable patterns of anxiety-related avoidance.

**Fig. 2.**
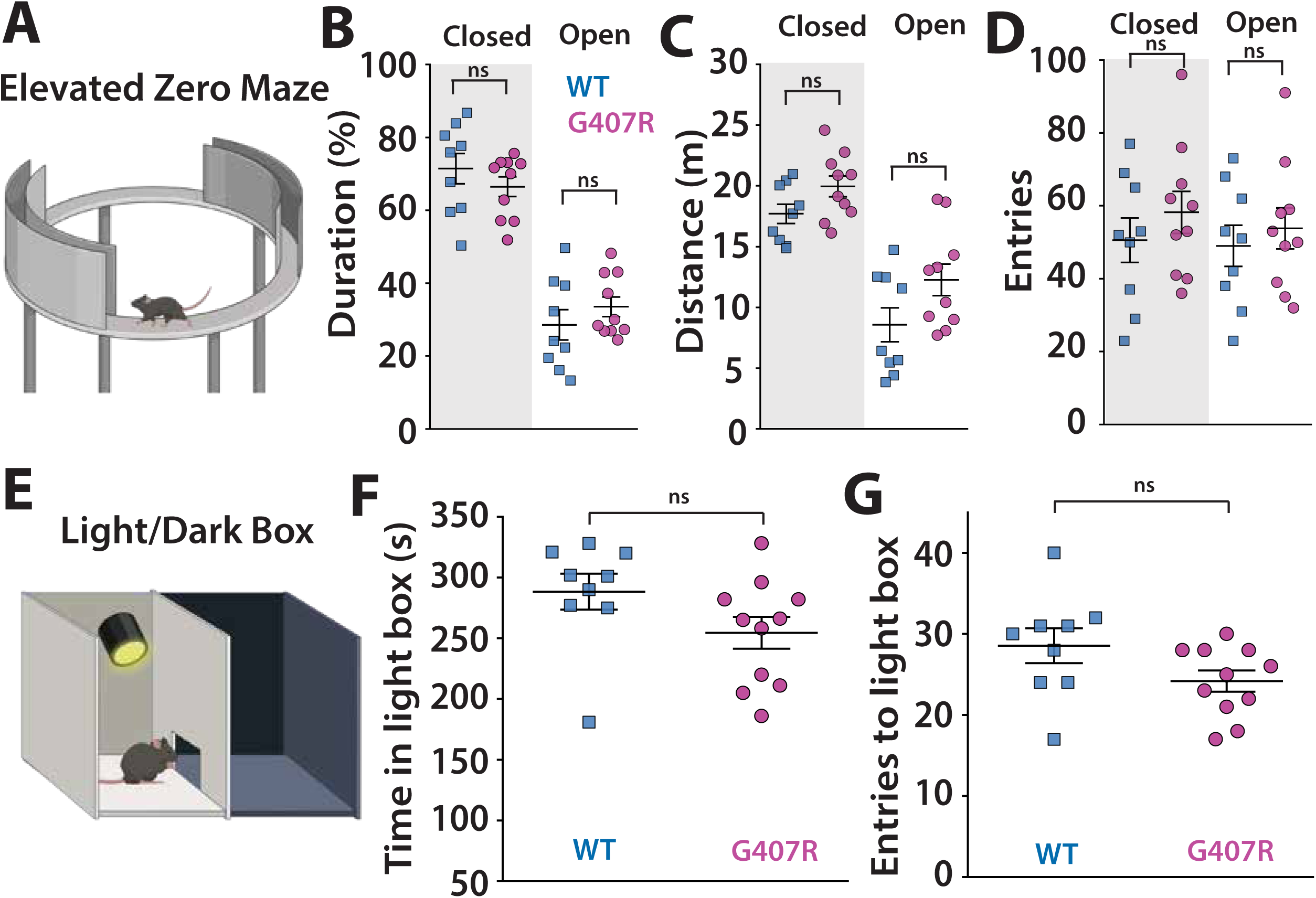
*Cacna1d*^G407R^ heterozygous mice exhibited normal anxiety-like behavior. (**A**) Performance of female *Cacna1d*^G407R^ heterozygous mice in the elevated zero. Schematic of the apparatus. (**B**) *Cacna1d*^G407R^ heterozygous mice showed normal behavior compared spending similar percentage of time in closed and open areas (**C**), distance traveled in each quadrant and (**D**) number of entries into the quadrants. *n* = 9 (wild-type) and *n* = 10 (heterozygous).(**E**) Schematic of the light/dark box. (**F**) Time spent in the light compartment and (**G**) the number of entries into the light compartment did not differ between genotypes. *n* = 9 (wild-type) and *n* = 11 (heterozygous). Schematics were generated in bioRender.

Both genotypes spent substantially more time in the closed quadrants than in the open quadrants, reflecting the expected preference for protected spaces (LME main effect of Zone, p < 0.0001; main effect of Genotype, p = 0.32; interaction, p = 0.15). Distance traveled in each quadrant showed the same pattern, with all mice moving more within the closed quadrants and no overall difference between genotypes (Zone p < 0.0001; Genotype p = 0.16; interaction p = 0.35). Numbers of entries into the quadrants were also similar between WT and mutant mice, with a slight reduction in open-quadrant entries consistent with avoidance of exposed areas (main effect of Zone, p < 0.0001; main effect of Genotype, p = 0.35). Together, these analyses indicate that zero-maze performance is broadly similar between WT and mutant mice, with no evidence for altered anxiety-like behavior in this assay.

As another test for anxiety-like behavior, we evaluated mice in the light/dark box, an assay that leverages the natural conflict between a rodent’s aversion to brightly illuminated, exposed environments and its drive to explore novel spaces (Crawley and Goodwin, 1980; Bourin and Hascoet, 2003)(**Figure 2E**). WT and G407R mutant mice spent similar amounts of time in the light compartment, and no significant genotype difference was detected (p = 0.074, Mann–Whitney U test)(**Figure 2F**). The number of entries into the light compartment was also comparable across groups, indicating similar exploratory drive and transitions between chambers (p = 0.087, Student’s t-test)(**Figure 2G**). Taken together, results from the open field, elevated zero maze, and light/dark box demonstrate that G407R mutant mice exhibit normal anxiety-like behavior across multiple relevant assays, with no significant genotype differences detected in any measure.

### Sociability, Social novelty and Social memory

Because social communication is often altered in human neurodevelopmental disorders, we evaluated social behavior in the *Cacana1d*^G407R^ mouse model using the three-chamber assay. The sociability phase provides a robust, widely validated measure of spontaneous social approach, with typical mice preferring interaction with an unfamiliar conspecific over an empty chamber (Moy et al., 2004; Silverman et al., 2010). Although the subsequent social novelty phase is generally considered less sensitive and may not reliably distinguish subtle genotypic differences, it remains a standard assay for assessing short-term social recognition memory supported by hippocampal and cortical circuits. We therefore tested mice in the three-chamber assay as we have previously described (Xu et al., 2021).

In the habituation phase with two identical empty cups, wild-type (WT) and mutant mice showed no evidence of side bias. Time spent at the left versus right empty cup did not differ within either genotype (WT: 115.1 ± 10.0 s vs 121.6 ± 9.6 s, p = 0.738, paired t-test; G407R: 125.7 ± 8.5 s vs 112.3 ± 8.8 s, p = 0.429, paired t-test)(**Figure 3A-C**). A discrimination ratio (see methods) was calculated to quantify potential left–right preference. Ratios were not different from zero in either genotype (WT: –0.011 ± 0.032, p = 0.738, one-sample t-test; G407R: 0.022 ± 0.027, p = 0.429, one-sample t-test) and did not differ between genotypes (p = 0.433, Welch’s t-test), indicating that baseline exploration of the chambers was balanced and comparable across groups (**Figure 3D**).

**Fig. 3.**
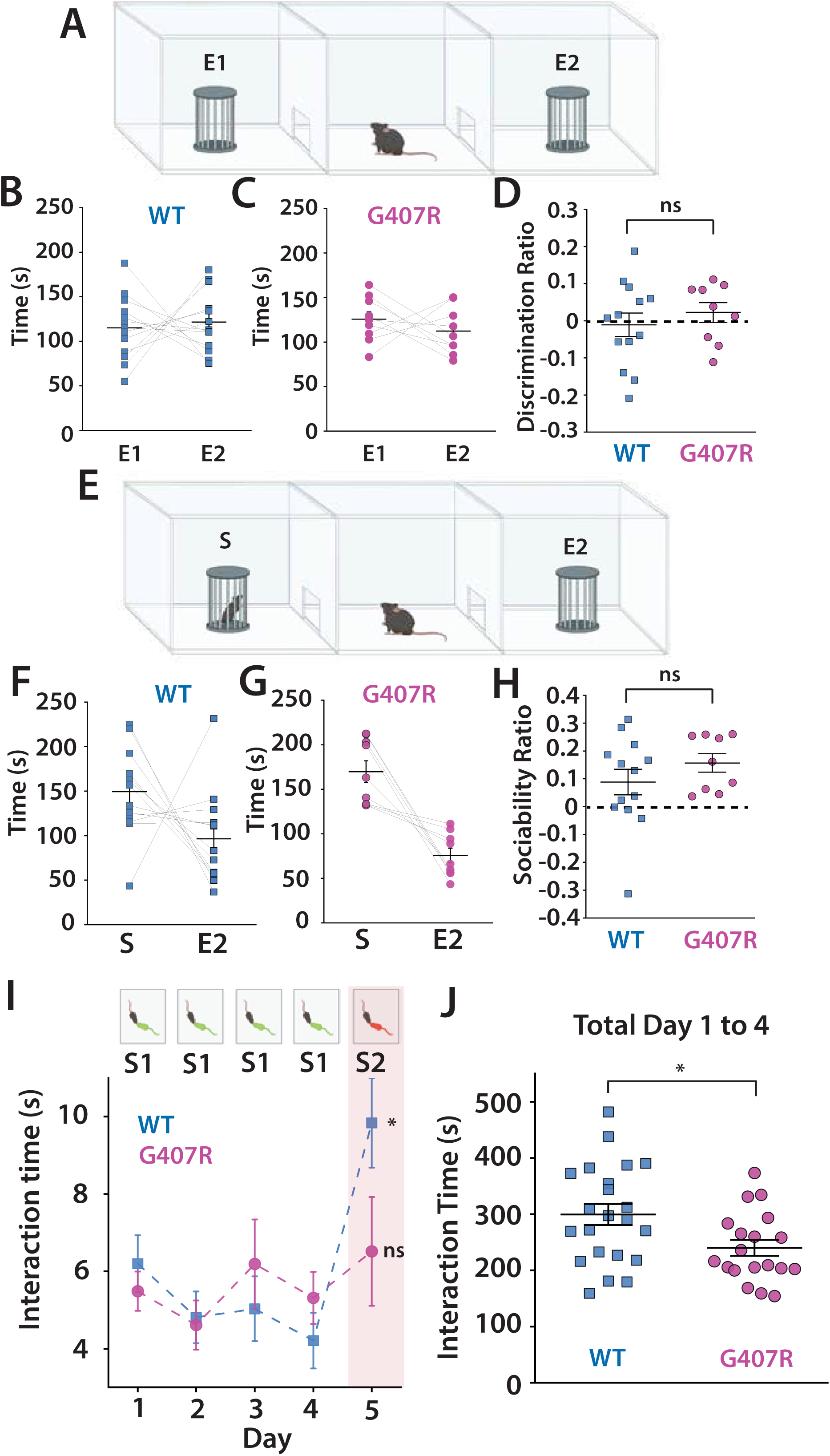
Impairment of social memory in *Cacna1d*^G407R^ heterozygous mice. (**A**) Schematic of the habituation phase in the 3-chamber test. (**B**) Time spent investigating the left and right cages by wild-type and (**C**) *Cacna1d*^G407R^ heterozygous mice. (**D**) The discrimination ratio reflecting left-right investigating bias did not differ between genotypes. (**E**) Schematic of the sociability phase in the 3-chamber test. (**F**) Duration of investigating a stranger conspecific (S) versus an empty cage (E2) by wild-type and (**G**) *Cacna1d*^G407R^ heterozygous mice (**H**) The ratio of time spent investigating the stranger animal relative to the empty cage did not differ between genotypes. *n* = 13 (wild-type) and *n* = 9 (heterozygous).(**I**) Schematic of the five-day social recognition/novelty test (top) and duration of contact initiated by the test mouse toward stranger 1 (S1) and stranger 2 (S2) across session (bottom panel). (**J**) Total interaction time during day 1-4. *n* = 22 (wild-type) and *n* = 19 (heterozygous). Schematics were generated in bioRender.

In the sociability phase, where one cup contained a novel conspecific and the other remained empty (Figure 3E), WT mice spent more time with the social stimulus than the empty cup (149.5 ± 13.5 s vs 96.5 ± 14.5 s; t(12) = 1.92, p = 0.079, paired t-test), although this difference did not reach statistical significance (Figure 3F). Mutant mice showed a clear and significant preference for the social stimulus (169.8 ± 12.2 s vs 75.7 ± 8.2 s; W = 0, p = 0.0039, Wilcoxon signed-rank test) (Figure 3G). A sociability index (see methods) was calculated for each animal and was not different between genotypes (WT: 0.088 ± 0.046; G407R: 0.157 ± 0.033; U = 42.0, p = 0.285, Mann–Whitney U test)(**Figure 3H**). Thus, both WT and mutant mice exhibit intact social approach behavior in the three-chamber assay, with no significant genotype effect on the overall strength of sociability.

Because the three-chamber assay primarily probes acute social behavior with single-trial social approach, we next asked whether the mutant mice can form stable social recognition across repeated encounters. To do this, we used a 5-day social recognition / social novelty paradigm in which mice were re-exposed to the same stimulus animal for four consecutive days and then presented with a novel conspecific on day 5. Similar multi-trial social habituation-dishabituation designs are widely used to study social recognition memory and its circuit mechanisms, with repeated encounters producing a progressive decline in investigation of the familiar mouse and a recovery of investigation when a novel mouse is introduced (Wang and Zhan, 2022). We quantified both the first 2 min of each 15-min session, when social investigation is highest and most sensitive to familiarity, and the total 15-min investigation time (Wang and Zhan, 2022). In each case we report the active investigation by the subject mouse of the stranger. WT mice showed a gradual decline in investigation of the repeated stimulus mouse across Days 1–4 (Day 1: 6.20 ± 0.74 s vs Day 4: 4.21 ± 0.72 s, Wilcoxon signed rank p = 0.019), consistent with intact social habituation (**Figure 3I**). In contrast, G407R mutant mice did not show any significant change in contact duration across Days 1–4 (Day 1: 5.49 ± 0.51 s vs Day 4: 5.31 ± 0.67 s, Wilcoxon signed rank, p = 0.89) (*Figure 3I*), When total social interaction across the full 15-min sessions (Days 1–4) was analyzed, mutant mice exhibited significantly reduced interaction time compared with WT controls (WT: 299.27 ± 18.59 s; Mutant: 239.89 ± 14.14 s (p = 0.015, Welch’s t-test)(**Figure 3J**).

To test social novelty recognition a novel mouse S2 was introduced on day 5 (S2)(**Figure 3I**). We compared investigation during the final familiar exposure (Day 4) with investigation of the novel conspecific S2 on Day 5 using the first 2 min of each session. WT mice showed a robust novelty response, with a significant increase in contact duration on Day 5 compared with Day 4 (Day 4: 4.21 ± 0.72 s; Day 5: 9.84 ± 1.16 s; p = 0.00027, paired t-test). In contrast, mutant mice failed to show a significant increase in investigation with the novel mouse (Day 4: 5.31 ± 0.67 s; Day 5: 6.51 ± 1.41 s; p = 0.651, Wilcoxon signed-rank test). These results indicate that while WT mice exhibit intact social novelty recognition, this behavior is impaired in G407R mutant mice.

### Automated spatial discrimination and reversal task to test behavioral flexibility

Our initial assessment of these mice had demonstrated that they appear to have behaviors consistent with elevated perseveration including digging and grooming behaviors. As these are key features of executive dysfunction and compulsive behavior prevalent in neuropsychiatric disease, we assessed behavioral flexibility and perseverative responding more rigorously by testing mice in an automated touch screen visual spatial discrimination and reversal learning task. Reversal learning assays are widely used to probe the ability to update stimulus–response associations when previously rewarded contingencies change, and increased errors following reversal are interpreted as a measure of perseveration and cognitive inflexibility (Ragozzino, 2007; Bissonette and Powell, 2012). Critically, the transition from goal-directed to habitual responding during extended training is mediated by striatal circuitry, with early learning relying on dorsomedial striatum (DMS) and overtraining recruiting dorsolateral striatum (DLS), which promotes rigid, habit-like stimulus–response behavior (Graybiel, 2008; Yin et al., 2009). Disruption of these striatal circuits leads to abnormal perseveration and impaired reversal learning, making this task a sensitive probe of striatal-dependent behavioral control.

We implemented this assay using an automated touchscreen operant conditioning system based on our previously developed open-source platform, Operant House (Otsuka, 2025). The apparatus consists of two touch-sensitive panels, one designated as the rewarded stimulus (S+) and the other as the unrewarded stimulus (S−) (**Figure 4A**). During the acquisition phase, touching the correct panel resulted in delivery of a water reward, whereas responses at the incorrect panel were not reinforced (Figure 4A). Stimulus positions remained fixed during acquisition and subsequent overtraining and were reversed at the onset of the reversal phase such that the previously unrewarded panel became rewarded (Figure 4A). To parametrically manipulate the strength of habit formation and the degree of perseverative responding, mice underwent short (2 days), intermediate (7 days), or long (14 days) overtraining after first reaching a performance criterion of ≥80% correct responses during acquisition, an approach shown to progressively bias behavior toward striatal-dependent habitual control.

**Fig. 4.**
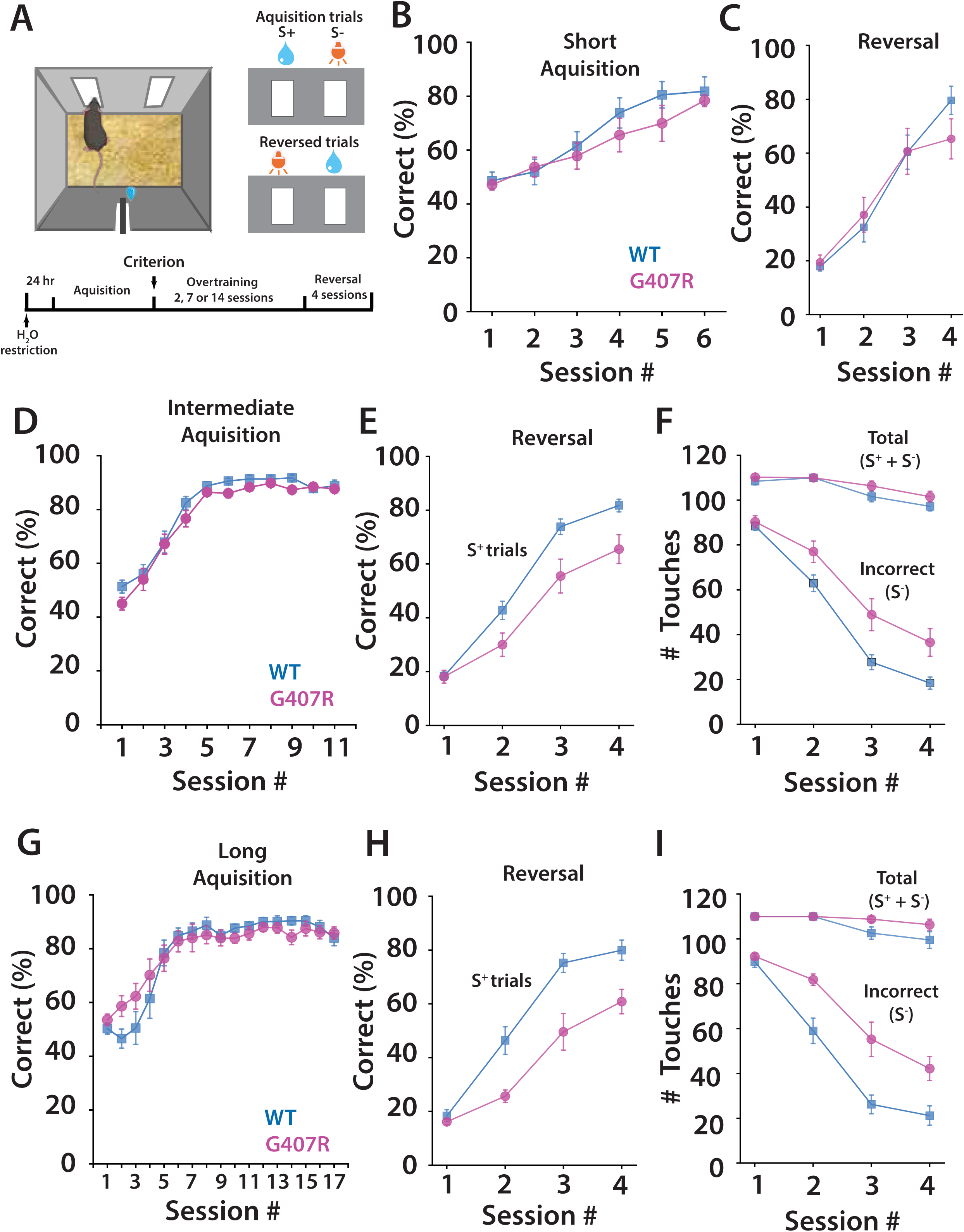
*Cacna1d*^G407R^ heterozygous mice showed perseverative behavior in the spatial discrimination task. (**A**) Schematic of the operant chamber and the task schedule. (**B**) Results from short (2-day) overtraining paradigm. Correct response rate in each session during the first 6 days of the acquisition phase and (**C**) the reversal phase in *Cacna1d*^G407R^ heterozygous and wild-type mice. (**D**) Performance in the intermediate (7-day) overtraining paradigm. Correct response rate during the first 11 days of the acquisition phase and (**E**) the reversal phase. (**F**) Total and incorrect touch responses across reversal sessions. (**G**) Results from the long (14-day) overtraining paradigm. Correct response rate during the first 17 days of the acquisition phase and (**H**) the reversal phase. (**I**) Total and incorrect touch responses across reversal sessions. Short overtraining: *n* = 10 (wild-type) and *n* = 10 (heterozygous); intermediate overtraining: *n* = 17 (wild-type) and *n* = 18 (heterozygous); Long overtraining: *n* = 10 (wild-type) and *n* = 10 (heterozygous)

During initial acquisition under the short (2-day) overtraining condition, both WT and mutant mice exhibited robust learning across sessions, as indicated by a strong main effect of session (F(5,84) = 40.99, p = 3.8 × 10⁻²¹)(**Figure 4B**). Moreover, WT and mutant mice were not significantly different in their performance during the acquisition phase, indicating that the two groups learned at comparable rates (**Figure 4B**). During reversal, both groups again showed strong improvement across sessions (main effect of session: F(3,57) = 76.14, p = 6.3 × 10⁻² ), and a significant main effect of genotype persisted (F(1,57) = 5.30, p = 0.025), with WT mice achieving higher late reversal performance (**Figure 4C**). The genotype × session interaction did not reach significance (F(3,57) = 2.30, p = 0.086), indicating that mutant mice were not fundamentally impaired in reversal learning rate, but instead exhibited a stable downward shift in performance across sessions.

In a second cohort of mice we elongated training 7 days past the 4-day acquisition (intermediate acquisition). Across the acquisition phase of the spatial discrimination task, both WT and G407R mutant mice showed robust learning curves and steadily increasing accuracy over sessions. A two-way repeated-measures ANOVA revealed a significant main effect of session on performance (*p* < 0.0001, RM-ANOVA), confirming that both groups learned the discrimination rule. However, there was no main effect of genotype on acquisition accuracy (*p* > 0.05, RM-ANOVA), nor a genotype × session interaction (*p* > 0.05, RM-ANOVA). Thus, the mutant mice acquired the initial stimulus–response rule at a rate comparable to WT controls. These results demonstrate that the basic instrumental learning and discrimination capabilities of the mutant mice remain intact prior to reversal training (**Figure 4D**).

When the rewarded and unrewarded positions were swapped (reversal), WT and G407R mutant mice showed markedly different behavioral profiles. WT mice exhibited an expected drop in accuracy early in reversal followed by steady improvement, whereas mutant mice displayed a more prolonged period of errors consistent with greater perseveration. A two-way repeated-measures ANOVA across reversal sessions showed a significant main effect of session (*p* < 0.0001, RM-ANOVA), reflecting overall adjustment to the new rule. Importantly, there was a significant main effect of genotype (*p* < 0.05, RM-ANOVA) and a significant genotype × session interaction (*p* < 0.05, RM-ANOVA), indicating that mutant mice were specifically impaired in reversal learning.

Together, this pattern reflects reduced behavioral flexibility and greater reliance on the previously reinforced strategy in the mutant mice (**Figure 4E**).

Analysis of the number of incorrect responses (S^-^) differed significantly between groups. A two-way repeated-measures ANOVA a significant effect of genotype (*p* < 0.01, RM-ANOVA), with mutants making substantially more incorrect touches than WT mice.

Incorrect touches therefore provide an additional quantitative marker of perseverative responding, reinforcing the conclusion that mutant mice have selective difficulty inhibiting outdated action–outcome contingencies (**Figure 4F**). Therefore, G407R mutant mice display normal acquisition of an initial spatial discrimination but show robust deficits during reversal, characterized by elevated perseverative errors.

To assess whether extended overtraining altered the acquisition or flexibility of stimulus–response learning, in a further cohort of mice we performed a long acquisition overtraining mice for 14 sessions after they initially reached a >80% correct rate in training. During the acquisition phase, both WT and G407R mutant mice showed the expected improvement in accuracy across sessions. Repeated-measures ANOVA revealed a significant effect of session in WT mice (RM-ANOVA: *F*(16,144) = 52.31, *p* < 0.0001) and in mutant mice (RM-ANOVA: *F*(16,128) = 31.57, *p* < 0.0001), indicating successful learning in both groups (**Figure 4G**). Direct genotype comparisons across acquisition sessions showed no significant WT vs mutant differences (*p* > 0.05, independent-samples tests), demonstrating that basic discrimination learning and response–reward mapping were intact in mutant animals. Thus, neither motivational nor associative processes required for initial task acquisition were impaired, even under extended training conditions (**Figure 4G**).

Following long overtraining, mice were shifted to reversal learning, in which the previously incorrect touchscreen location became rewarded. WT mice showed the expected rapid reorganization of responding, with accuracy increasing across reversal sessions (RM-ANOVA: *F*(3,27) = 29.44, *p* < 0.0001). Mutant mice also demonstrated a significant effect of session (RM-ANOVA: *F*(3,24) = 12.89, *p* < 0.001), indicating that they were able to eventually acquire the new contingency. However, the pattern of performance differed markedly between genotypes: WT and mutant accuracy did not differ on the first reversal session (Day 1; *p* > 0.1), but mutants performed significantly worse on all subsequent sessions (Day 2: *p* < 0.01; Day 3: *p* < 0.001; Day 4: *p* < 0.01; independent-samples tests)(**Figure 4H**). These findings indicate that mutant mice exhibit a pronounced reduction in behavioral flexibility, with a strong tendency to perseverate on the previously learned (but now incorrect) response strategy.

To distinguish flexibility impairments from possible differences in task engagement, motivation, or motor output, we quantified total touchscreen responses and incorrect touches during reversal. Total touches (S^+^ + S^-^) did not differ between genotypes on any session (all *p* > 0.10, independent-samples tests), indicating that both groups were comparably engaged in the task and capable of performing the required motor actions (**Figure 4I**). In contrast, incorrect touches (S^-^) revealed a robust genotype effect (**Figure 4I**). WT mice displayed a strong, session-dependent reduction in errors (RM-ANOVA: *F*(3,27) = 35.41, *p* < 0.0001), while mutants also showed a session effect (RM-ANOVA: *F*(3,24) = 14.72, *p* < 0.0001) but maintained persistently higher error rates. Between-genotype comparisons showed no difference on Day 1 (*p* > 0.1), but mutants committed significantly more errors on Days 2–4 (Day 2: *p* < 0.0001; Day 3: *p* < 0.001; Day 4: *p* < 0.01)(**Figure 4I**). These elevated error rates, in the absence of reduced total touches, demonstrate that mutants specifically fail to suppress the previously reinforced response, consistent with enhanced perseveration rather than reduced motivation or motor impairment.

Taken together, these results demonstrate that while acquisition learning remains intact in mutant mice even under extended training conditions that promote habit formation, the same animals show a marked deficit during reversal learning, characterized by impaired suppression of perseverative responses and reduced behavioral flexibility.

This pattern suggests a selective disruption in cortico-striatal mechanisms supporting the transition from habitual responding to updated, goal-directed control.

To further probe cognitive flexibility under conditions of uncertainty, we employed an intermediate acquisition protocol followed by probabilistic reversal learning (PRL) task in which the newly correct response was rewarded on 80% of trials and punished on 20%, while the previously correct (now incorrect) response was reinforced on 20% of trials and punished on 80% (**Figure 5A**). Probabilistic feedback structures are widely used in both rodents and humans to address the involvement of prefrontal–striatal circuits involved in updating stimulus–outcome associations, evaluating action outcomes under uncertainty, and suppressing perseverative responding (Dalton et al., 2016). Unlike deterministic reversal tasks, probabilistic reversal requires animals to integrate feedback across multiple trials and tolerate misleading outcomes, making it especially sensitive to disruptions in orbitofrontal cortex (OFC) function and striatal gating (Conn et al., 2025; Koloski et al., 2025). Prior studies have shown that PRL paradigms reveal deficits not detected in deterministic reversal learning, particularly in models of neuropsychiatric conditions involving compulsivity, autism-related perseveration, or corticostriatal dysregulation (Amodeo et al., 2012). Thus, introducing uncertainty through probabilistic reinforcement allows us to assess higher-order cognitive flexibility, including sensitivity to feedback, ability to shift behavioral strategies, and resilience against habitual responding, domains highly relevant to striatal-dependent learning and the phenotype observed in the G407R mutant mice.

**Fig. 5.**
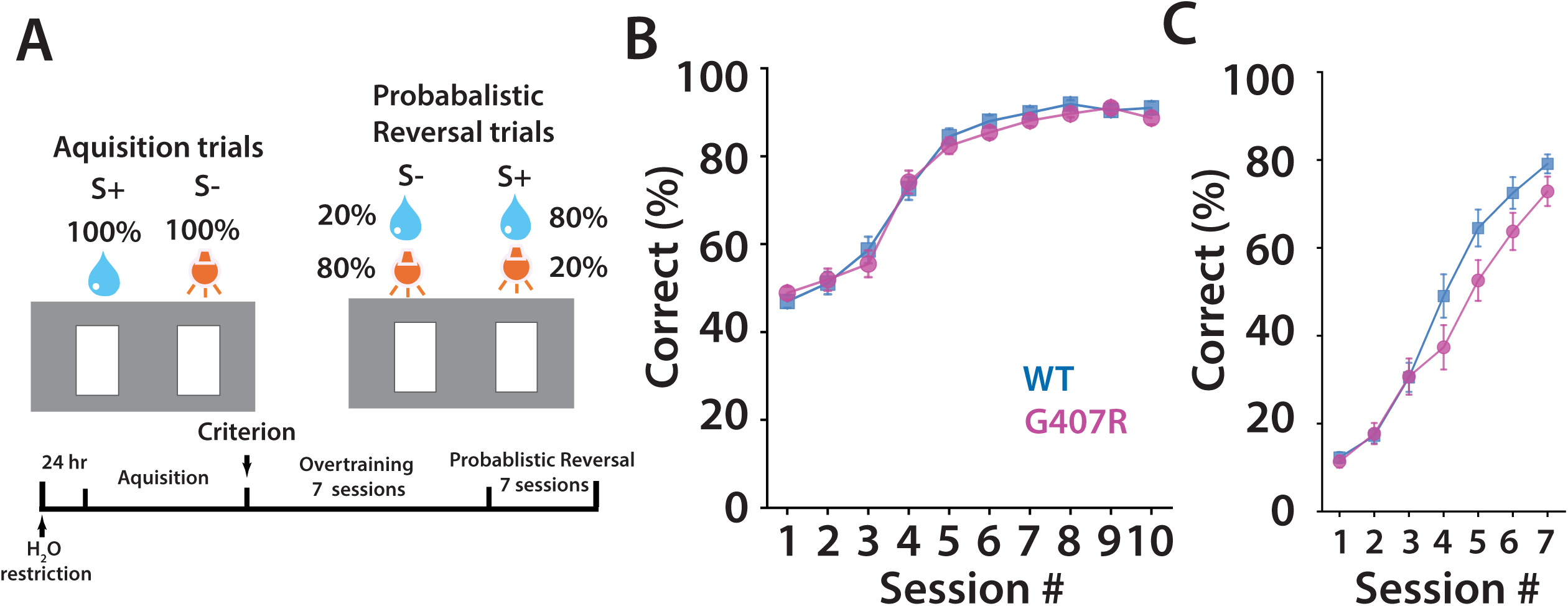
*Cacna1d*^G407R^ heterozygous mice demonstrate perseverative phenotype in the probabilistic spatial discrimination task. (**A**) Schematic of task structure and rule for the acquisition and reversal in the probabilistic spatial discrimination task. (**B**) Mean correct response rate during acquisition phase and (C) reversal phase *n* = 27 (wild-type) and *n* = 25 (heterozygous).

Mice were first trained and performance improved robustly across acquisition sessions in both genotypes, reflected by a significant main effect of session (two-way repeated-measures ANOVA, *p* < 0.0001)(**Figure 5B**). There was no significant main effect of genotype (n.s.), indicating that overall acquisition performance did not differ between WT and mutant mice. Moreover, there was no significant session × genotype interaction (n.s.), demonstrating that the rate and trajectory of learning across sessions were comparable between groups. Together, these results indicate that as before basic acquisition is preserved in G407R mutant mice, and that learning deficits are not evident during initial task acquisition (**Figure 5B**)

Performance during probabilistic reversal improved across sessions in both WT and G407R mutant mice, reflected by a robust main effect of session (two-way repeated-measures ANOVA, *p* < 0.0001)(**Figure 5C**). There was no significant main effect of genotype on overall performance (*p* > 0.1), indicating that both groups were able to acquire the reversed contingency under probabilistic reinforcement. However, a significant session × genotype interaction was observed (RM-ANOVA, *p* < 0.05), indicating divergent learning trajectories across reversal sessions (**Figure 5C**). Post hoc comparisons revealed that WT mice showed a steady and significant increase in correct responding across later reversal sessions (*p* < 0.01), whereas mutant mice exhibited slower improvement and lower performance during intermediate reversal sessions (*p* < 0.05, genotype comparison), consistent with increased perseverative responding under conditions of outcome uncertainty. By the final reversal sessions, performance converged between genotypes (*p* > 0.1), suggesting that mutant mice retain the capacity to update stimulus–outcome associations but require additional experience when reinforcement contingencies are probabilistic (**Figure 5C**).

## Discussion

Restricted and repetitive behaviors, including behavioral inflexibility and perseveration, are a core diagnostic domain of autism spectrum disorder (ASD) and are strongly associated with impairments in adaptive functioning and cognitive control (Granader et al., 2014). Perseverative behaviors manifest across multiple levels in ASD, ranging from repetitive motor actions to rigid cognitive strategies and resistance to change.

Importantly, these features are not only clinically salient but are thought to reflect disruptions in corticostriatal circuits that govern action selection, habit formation, and flexible updating of behavioral rules (Graybiel, 2008). Consistent with this, a wide range of ASD-associated mouse models, including mutations in *Shank3*, *Fmr1*, and *Cntnap2*, exhibit perseverative responding and impaired reversal learning supporting the translational relevance of these behaviors as measurable circuit-level endophenotypes (Peca et al., 2011; Penagarikano et al., 2011; Mercaldo et al., 2023).

In the present study, we identify heightened perseveration and impaired behavioral flexibility in *Cacna1d*^G407R^ mutant mice across multiple, independent behavioral assays. While basic anxiety-like behaviors and general locomotor activity were largely preserved, G407R mutants consistently showed deficits specifically in tasks that required updating previously learned action–outcome associations or disengaging from over-trained response strategies. In spatial discrimination paradigms, both genotypes acquired the initial rule efficiently, indicating intact basic learning and motivation.

However, following extended training, G407R mutant mice exhibited increased perseveration during deterministic reversal, characterized by prolonged responding at the previously rewarded location, elevated incorrect responses, and delayed acquisition of the new rule. These effects scaled with training history and were most pronounced after long overtraining, a condition known to bias control toward dorsolateral striatal habit systems.

Strikingly, perseverative phenotypes in G407R mutants generalized across task structure and reinforcement contingencies. In probabilistic reversal learning where ambiguity and stochastic feedback place additional demands on cognitive flexibility, G407R mutant mice again showed impaired adaptation, increased incorrect responding, and reduced sensitivity to changing contingencies. This pattern suggests that the observed deficits are not limited to deterministic rule switching, but extend to environments requiring integration of uncertain feedback, a process heavily dependent on frontostriatal and orbitofrontal circuitry. Together, these findings demonstrate that the G407R mutation disrupts mechanisms that constrain habitual responding and enable flexible behavioral updating, rather than broadly impairing learning, motivation, or affective state.

Importantly, perseverative tendencies were also evident in ethologically distinct assays, including nestlet shredding and digging, reinforcing the interpretation that *Cacna1d*^G407R^ mutant mice exhibit a convergent inflexibility phenotype rather than task-specific artifacts. The consistency of these effects across instrumental, probabilistic, and naturalistic behaviors strengthens the conclusion that altered perseveration represents a core behavioral consequence of the *Cacna1d*^G407R^ mutation. Taken together, our results align G407R mutant mice with a growing class of ASD-relevant models in which exaggerated habit formation and impaired reversal learning that likely reflect disrupted corticostriatal control, providing a mechanistic entry point for linking molecular dysfunction to circuit-level rigidity and behavioral inflexibility.

### Cacna1d^G407R^mice and circuit mechanisms underlying behavioral flexibility

*De novo* gain-of-function mutations in CACNA1D, which encodes the L-type calcium channel Cav1.3, have been repeatedly identified in individuals with autism spectrum disorder and related neurodevelopmental phenotypes, often accompanied by motor stereotypies, cognitive rigidity, and altered reward processing (Pinggera et al., 2015; Pinggera et al., 2017; Dannenberg et al., 2024). Among these, the G407R variant is one of the best-characterized pathogenic alleles and produces a pronounced hyperpolarizing shift in channel activation, increased window current, and enhanced calcium influx at physiologically relevant membrane potentials (Pinggera et al., 2018).

Cav1.3 channels are highly expressed in brain regions implicated in behavioral flexibility, including the dorsal and ventral striatum, prefrontal cortex, and basal ganglia output nuclei, where they play a critical role in activity-dependent calcium signaling and synaptic plasticity (Striessnig et al., 2006; Kabir et al., 2017). In striatal medium spiny neurons, L-type calcium channels, particularly Cav1.3, are essential for plasticity at corticostriatal synapses, processes that are central to updating action–outcome contingencies and suppressing previously reinforced behaviors (Adermark and Lovinger, 2007; Rafalovich et al., 2015; Zhai et al., 2024).

Disruption of striatal plasticity has been demonstrated to bias behavior toward habitual and perseverative responding, reducing behavioral flexibility during reversal and probabilistic learning tasks (Shan et al., 2015). The *Cacna1d*^G407R^ mice have elevated Ca^2+^ influx through L-type channels in striatal projection neurons which disrupts a form of long term depression (Zhai et al., 2024) impairing the normal re-weighting of corticostriatal circuits required for flexible behavior.

Together, these findings position the *Cacna1d*^G407R^ mutation as a biologically plausible driver of perseveration through dysregulated calcium-dependent plasticity in corticostriatal circuits, providing a mechanistic framework for interpreting the robust deficits in reversal learning and behavioral flexibility observed across multiple assays in the present study.

In summary, the *Cacna1d*^G407R^ mutation produces a convergent phenotype characterized by exaggerated behavioral persistence and impaired flexibility across multiple assays. Mutant mice learned task contingencies normally but showed difficulty disengaging from previously reinforced strategies when contingencies changed, particularly following extended training and under probabilistic reinforcement. This inflexible behavioral profile was mirrored by increased repetitive behaviors in ethologically relevant assays, supporting a generalized shift toward perseverative action selection rather than a task-specific learning deficit. Together, these findings identify Cav1.3 gain-of-function as a mechanism that stabilizes behavioral states at the expense of adaptive updating, providing a mechanistic link between L-type calcium signaling, corticostriatal plasticity, and core features of autism related to behavioral rigidity.

## Methods

### Procedures

All procedures relating to animal care and treatment were performed according to the ethical policies and protocols approved by the Northwestern University IACUC. Behavioral tests were conducted during the light phase of the circadian cycle except for the nestlet shredding task, and were performed after at least 30-min acclimation period in the behavioral testing room.

### Animals

The *Cacna1d*^G407R^ mutation was engineered into the mouse genome as we previously described (Zhai et al., 2024). Briefly, a CRISPR/Cas9 gene editing system was used to substitute glycine at position 407 for an arginine. A <u>GG</u> to <u>AG</u> mutation was introduced in exon 8. Single guide RNAs (gRNA), Cas9 mRNA and a single strand oligonucleotide donor (ss donor) with a single base mutation (G407 to R407) were microinjected into mouse zygotes. Two female founders with the positively identified G407R mutation were bred with wild type C57/bl6J mice (Jackson laboratory), and each of these founders produced viable offspring with the G407R mutation. Heterozygous G407R mice were crossed with wild-type mice to generate G407R heterozygous and wild-type littermate controls for behavioral testing.

### Open-field test

Animals were placed in the middle of the arena (60cm × 60cm x 30 cm) at the start of the test. Mice were continuously tracked by an overhead video camera for 30 min and their positions tracked using EthoVision XT. The center region was defined as an area that covered 44.4 % of total arena. Data were analyzed using a linear mixed-effects model (the mixed-model analogue of a repeated-measures ANOVA), with time as a within-subject factor and genotype as a between-subject factor.

### Marble Burying Test

Tests were performed like what we have previously described (Xu et al., 2017). A standard cage (36 cm × 31 cm × 19 cm) with ∼3-5 cm of bedding had twenty-four dark marbles placed in six columns of four. Animals were introduced to this new cage and remained inside for 15 minutes. Experiments were video recorded with cameras positioned above the cage and animal’s cumulative time spent digging was scored post hoc. After 15 minutes mice were removed from the cage and the number of marbles that were buried by more than two-thirds of their diameter were measured without further perturbation of the cage or bedding.

### Induced Grooming

Spray-evoked grooming was assessed using a water-mist induction procedure. Each animal was placed individually into a clean, empty polycarbonate cage (30 × 20 × 15 cm) with no bedding or enrichment, and allowed to acclimate. A fine water mist was applied to the forehead using a handheld spray bottle delivering a single light spray sufficient to dampen the fur without startling the animal. Immediately after spraying, mice were video-recorded and total grooming duration was quantified offline by an experimenter blind to genotype. Grooming was defined according to established ethological criteria as coordinated facial, head, or body grooming movements, including paw strokes, licking, scratching, or fur manipulation. Pauses without grooming were scored as interruptions and not counted toward total grooming time. Between trials, cages were thoroughly cleaned with 70% ethanol to remove olfactory cues.

### Nestlet Shredding

Nestlet shredding was assessed as a measure of repetitive and compulsive-like behavior. Mice were singly housed in clean cages containing standard bedding and provided with a single pre-weighed cotton nestlet (∼2.5 g) at the onset of the testing period. Animals had ad libitum access to food and water throughout the assay. After leaving the animals for 12 hours, the remaining intact nestlet material was collected and weighed. Shredding was quantified as the percentage of the original nestlet mass that had been torn or fragmented.

### Water-Seeking Perseveration Task

Male mice that had previously undergone a spatial discrimination task were used in this experiment. Mice were individually housed in their homecage with standard wood-chip bedding, *ad libitum* food pellets and a water bottle for 8 days. The water bottle was then removed and food pellets were scattered on the cage floor to induce water deprivation for 30 hours. Following that, cages were transferred to the behavioral testing room under 20-lux illumination. An empty water bottle was placed on each cage and licking duration was recorded for 10 min.

### Zero maze test

The zero maze consisted of an elevated circular gray runway (outer diameter 56 cm, 5.5 cm width, raised 33 cm above the floor) separated into four quadrants with two quadrants on opposing sides protected by tall walls (14 cm high) and two quadrants remaining open. The maze was illuminated at 300 lux inside a sound-proof testing chamber. Mice were individually placed on an enclosed part of the runway and the mouse’s trajectory was tracked by an overhead video camera during a 10-minute session. Animal position was extracted using Mr. Behavior tracking software.

### Light/dark test

The apparatus consisted of two-compartments separated by an opening. Both the light and dark compartments were of the same dimensions (26(d) x 20(w) x 30(h) cm). The light chamber was illuminated with a fluorescent lamp (3500 lux) and the other dark box had no illumination (2.7 lux). The two compartments were separated by a wall with an opening (8.5 (w) x8(h) cm). Mice were placed in the center of the light chamber and recorded for 5 min using an overhead camera in the light chamber.

### Three Chamber Sociability test

Three-chamber social approach testing was performed in a standard Plexiglas apparatus consisting of three interconnected compartments of equal dimensions. The test was conducted under low ambient lighting. Stimulus animals were age- and sex-matched conspecifics that had no prior physical contact with the subject mice. All trials were recorded and scored using automated tracking software, with manual confirmation when necessary. Habituation Phase: At the beginning of the assay, the subject mouse was placed into the center chamber and allowed to freely explore all three chambers for 10 min. Two identical inverted wire cups were positioned on the left and right side chambers. No stimulus mouse was present during this phase. Time spent investigating each empty cup (E1 and E2) was quantified as the cumulative duration during which the subject’s nose was oriented toward and within proximity of the cup. This phase was used to measure baseline exploratory behavior and detect potential side preferences. A discrimination ratio (E1 - E2)/600 was calculated to quantify left–right bias across the 10-min (600 s) habituation session. Sociability Phase: Immediately following habituation, the subject was briefly confined to the center chamber while a novel stimulus mouse was placed under one wire cup (social location, S) and the opposite empty cup remained unchanged (E2). The side containing the stimulus mouse was counterbalanced across animals and genotypes. The subject was then allowed to re-enter all chambers for 10 min. Time spent investigating the social cup (S) versus the empty cup (E2) was quantified. A sociability index (S – E2)/600 was calculated to measure preference for social interaction normalized to total session duration.

### Five-day social recognition/novelty test

Social recognition and habituation to a repeated conspecific were assessed using a five-day direct interaction paradigm (Bariselli et al., 2018). Adult test mice were individually placed in a neutral Plexiglas arena (26 × 40 cm) with clean bedding and allowed to freely interact for 15 min with a sex- and age-matched juvenile stimulus mouse (S1) that had no prior contact with the subject. The same stimulus mouse was used on days 1–4 to assess social habituation, and on day 5 a novel, age- and sex-matched stimulus mouse (S2) was introduced to probe social recognition of a new conspecific. Stimulus mice were housed separately from test mice throughout the experiment, and arenas were thoroughly cleaned between sessions to remove olfactory cues. Sessions were video-recorded and the duration of the physical contact including sniffing of the head or body, close following and paw or body contact initiated by the test mouse was measured offline by a custom software using both the first 2 min of each session (to capture peak exploratory drive) and the full 15-min interaction period. Trials with sustained aggressive behavior were excluded *a priori*.

This paradigm is widely used to quantify progressive reduction in investigation across repeated exposures and recovery of exploration to a novel conspecific as an index of social habituation and social recognition memory.

### Spatial Discrimination and Reversal Learning

Spatial discrimination and reversal learning were assessed using an automated touchscreen-based operant conditioning system called Operant House (Otsuka, 2025). Mice were water-restricted 24 hours prior to the initial session to motivate task engagement. The test chamber (14.5(d) x 22(w) x 21.5 (h) cm) had an LCD monitor covered with a mask having two apertures to display rectangles as the optical stimulus (3(w) x 6.5(h) cm). Nose-poking was detected by the infrared camera located on the top of the mask. The floor was covered with cage bedding and food pellets. The chamber was not illuminated except for punishment. During the acquisition phase, mice learned to discriminate between two spatially distinct touchscreen locations. One location was designated as correct and the other as incorrect. Touching the correct location resulted in automated movement of the water nozzle making it accessible. Nose-poke to the water nozzle was detected by an infrared camera and the nozzle was retracted 2 sec after the initiation of the water access.

Touching the incorrect location resulted in no reward and was punished with a 15-sec ceiling illumination (150 lux). Each session continued until the subject attained 80 correct responses or reached a total of 110 responses. If the animal failed to attain 80 correct responses, it received supplemental water to ensure that total water intake did not depend on task performance (e.g. if its correct response was 60, it was allowed access to water nozzle for (80-60) x2 = 40 sec). After the session, the subject was returned to its homecage.

The spatial contingencies were fixed for each mouse throughout the acquisition phase. Training continued until mice reached a performance criterion of ≥80% correct responses within a session.

Following criterion attainment, mice underwent additional overtraining to manipulate the strength of learned stimulus–response associations. Three overtraining conditions were used: short acquisition (2 days of overtraining), intermediate acquisition (7 days of overtraining), or long acquisition (14 days of overtraining). The session where the subject first achieved ≥80% correct was included in this overtraining period. If the correct rate fell below 80%, that session was not counted as overtraining. Overtraining sessions were identical in structure to acquisition sessions, with spatial contingencies unchanged. After the completion of the overtraining sessions, the reversal phase began. In the reversal phase, the reward contingencies were switched such that the previously correct touchscreen location became incorrect, and the previously incorrect location became correct. No explicit cue signaled the contingency change. Mice were tested across multiple reversal sessions, and performance was quantified as percent correct responses, number of total touchscreen responses, and number of incorrect responses per session. These measures were used to assess behavioral flexibility and perseverative responding following different levels of overtraining. Experimenters were blinded to genotype during testing and data analysis.

### Probabilistic Spatial Discrimination Reversal

Mice were water-restricted for 24 hours and then tested under identical motivational and environmental conditions. Mice were first trained on the spatial discrimination task described above until they reached a criterion of ≥80% correct responses within a session, followed by 7 days of overtraining with fixed spatial contingencies. During the probabilistic reversal phase, reward contingencies were switched such that the previously incorrect location became the preferred choice. Unlike the deterministic reversal, reinforcement was delivered in a probabilistic manner: responses at the newly correct location were rewarded on 80% of trials and unrewarded/punished on 20% of trials, whereas responses at the newly incorrect location were rewarded on 20% of trials and unrewarded/punished on 80% of trials. No explicit cue signaled the reversal or the probabilistic nature of reinforcement. Probabilistic contingencies remained constant throughout the reversal phase.

Behavioral performance was quantified across sessions as percent correct responses, total number of touchscreen responses, and number of incorrect responses. This task design was used to assess behavioral flexibility and sensitivity to probabilistic feedback under conditions in which reward contingencies are partially ambiguous. Experimenters were blinded to genotype during behavioral testing and data analysis.

## Data analysis and statistics

All statistical analyses were performed in Python using NumPy, SciPy, and Pingouin libraries with custom analysis scripts. For each dataset, assumptions of normality were assessed using the Shapiro–Wilk test, and equality of variance was evaluated using Levene’s test where applicable. The choice between parametric and non-parametric tests was based on these assumption checks.

Within-Subject Comparisons: For comparisons made within the same animals across conditions (e.g., EL vs ER during three-chamber habituation; SL vs ER during sociability; Day 4 vs Day 5 during social novelty), paired t-tests were used when the distribution of difference scores met normality assumptions. When difference scores deviated from normality, the Wilcoxon signed-rank test was applied. These tests were selected because they account for repeated measurements within the same subjects and maximize statistical power under appropriate distributional conditions.

Repeated-Measures Across Days: For multi-day social habituation (Days 1–4), where the same mice were repeatedly tested across four sessions, within-genotype habituation was analyzed using the Friedman repeated-measures test, a non-parametric alternative to repeated-measures ANOVA that does not require normality. Follow-up comparisons of specific time points (e.g., Day 1 vs Day 4) were performed using Wilcoxon signed-rank tests. This approach was selected to robustly detect monotonic changes in social interaction across repeated exposures.

Between-Genotype Comparisons: For independent genotype comparisons (e.g., discrimination ratios, sociability indices, total 15-min interaction times, grooming duration, digging time, marble burying, light/dark measures), Welch’s unpaired t-test was used when data met normality assumptions, as it does not assume equal variances between groups. When normality was violated, the Mann–Whitney U test was used as a non-parametric alternative.

Preference and Discrimination Indices: Derived preference measures, including the three-chamber discrimination ratio and the sociability index, were tested against zero using one-sample t-tests when normally distributed or Wilcoxon signed-rank tests otherwise. These tests determine whether a group exhibits a systematic preference beyond chance (zero).

Anxiety and Zone-Based Measures: For assays involving repeated measurements across spatial zones (e.g., open vs closed arms in the zero maze), data were analyzed using two-way repeated-measures ANOVA with Greenhouse–Geisser correction when parametric assumptions were met, allowing assessment of main effects of zone, genotype, and their interaction.

Outlier Analysis: Potential extreme values in normally distributed paired difference datasets (e.g., Δ = SL − ER) were examined using Grubbs’ test (α = 0.05, two-sided).

General Reporting Standards: All tests were two-tailed, with statistical significance defined as α = 0.05. Data are presented as mean ± SEM unless otherwise noted. Exact test statistics, degrees of freedom, and p-values are reported in the Results section.

Sample sizes for each comparison are provided in the figure legends.

## Acknowledgements

This work was supported by NIH/NIMH *R01MH099114* and SFARI pilot award to AC.

## Author Contributions

SO, JX, AC designed the experiments SO and JX performed experiments, analyzed data and wrote the paper. S.O. developed the touch screen automated task and produced code to operate the procedures.SO, JX and AC prepared the manuscript.

